# The impact of CSF-filled cavities on scalp EEG and its implications

**DOI:** 10.1101/2023.11.30.569158

**Authors:** Vitória Piai, Robert Oostenveld, Jan Mathijs Schoffelen, Maria-Carla Piastra

## Abstract

Previous studies have found EEG amplitude and scalp topography differences between neurotypical and neurological/neurosurgical groups, being interpreted at the cognitive level. However, these comparisons are invariably accompanied by anatomical changes. Critical to EEG are the so-called volume currents, which are affected by the spatial distribution of the different tissues in the head. We investigated the effect of CSF-filled cavities on simulated EEG scalp data. We simulated EEG scalp potentials for known sources using different volume conduction models: a reference model (i.e., unlesioned brain) and models with realistic CSF-filled cavities gradually increasing in size. We used this approach for a single source close or far from the CSF-lesion cavity, and for a scenario with a distributed configuration of sources (i.e., a “cognitive ERP effect”). Magnitude and topography error between the reference and lesion models were quantified. For the single-source simulation close to the lesion, the CSF-filled lesion modulated signal amplitude with more than 17% magnitude error, and topography with more than 9% topographical error. Negligible modulation was found for the single source far from the lesion. For the multi-source simulations of the cognitive effect, the CSF-filled lesion modulated signal amplitude with more than 6% magnitude error, and topography with more than 16% topography error in a non-monotonic fashion. In conclusion, the impact of a CSF-filled cavity cannot be neglected for scalp-level EEG data. Especially when group-level comparisons are made, any scalp-level attenuated, aberrant, or absent effects are difficult to interpret without considering the confounding effect of CSF.

**Impact statement:** Previous studies have found EEG amplitude and scalp topography differences between neurotypical and neurological/neurosurgical groups (whose brain damage leads to the presence of a CSF-filled cavity), being interpreted at the cognitive level. Via simulations of scalp-level EEG patterns, we show that attenuated, aberrant, or absent effects in these comparisons are difficult to interpret without considering the confounding effect of CSF.

**Funding:** This study was partly supported by grants from the Netherlands Organization for Scientific Research (Nederlandse Organisatie voor Wetenschappelijk Onderzoek [NWO]) to V. P. (451-17-003 and VI.Vidi.201.081) and to the Language in Interaction Consortium (024-001-006).

**CRediT:** **VP:** Conceptualization, Data curation, Formal Analysis, Funding acquisition, Investigation, Methodology, Project administration, Visualization, Writing – original draft, Writing – review & editing

**RO:** Conceptualization, Methodology, Software, Writing – review & editing

**JMS:** Conceptualization, Methodology, Software, Writing – review & editing

**MCP:** Conceptualization, Investigation, Methodology, Visualization, Writing – review & editing

## 1. Introduction

The electroencephalogram (EEG) – the recording of the electrophysiological signal over the scalp – reflects post-synaptic potentials of thousands of synchronously activated neurons (Lopes da Silva, 2013). Given its excellent temporal resolution and direct relationship with brain activity, the electrophysiological signal has played a pivotal role in our understanding of cognitive and sensorimotor functions, as well as changes in these functions following brain damage. Importantly, in certain cases of brain damage (e.g., stroke or surgical resection), not only functional but also structural brain changes may affect aspects of the measured electrophysiological signals, impacting the inferences one can draw from them.

A common type of brain damage studied with scalp electrophysiology is stroke. Stroke is caused either by the lack of blood flow to a region or by bleeding, with the resulting poor perfusion causing cell death. The area occupied by the dead tissue eventually becomes filled with cerebrospinal fluid (CSF), which is the most conductive element of the head. Other cases that lead to CSF-filled cavities are surgical resections, for example of a tumour or of pathological tissue causing epilepsy.

Various cognitive and sensorimotor domains have been studied with electrophysiology following brain damage, most commonly using event-related potentials (ERPs). For example, in the domain of acquired language disorder (i.e., aphasia) due to stroke, when comparing the stroke group with a stroke-free group, it has been found that ERP components tend to have smaller amplitudes over the scalp and unusual scalp distributions. Moreover, studies have also found that aspects of ERP components (e.g., amplitude and topography) are correlated with the severity of the stroke and the corresponding impairment (for reviews e.g., Ehlers et al., 2015; Meechan et al., 2021; Silkes & Anjum, 2021; Vatinno et al., 2022).

These seemingly functional changes are invariably accompanied by anatomical changes that need careful consideration. In particular, it is critical to consider the effects of CSF in these comparisons and the extent to which differences measured at the scalp can be (partially) trivially explained by underlying differences in the presence and spatial distribution of CSF in relation to the brain tissue. Impressed neuronal currents in the brain only indirectly result in electric potentials, as measured over the scalp with EEG. Crucial to the EEG signal are the so-called volume currents, which flow through the various tissues in the head (grey and white matter, CSF, skull, and scalp). The spatial distribution of the different elements will critically affect the path that the volume currents will take, and thus impact the EEG signal. In particular, it is necessary to consider the effects of the redistribution of CSF, given its high conductivity.

The biophysics literature has provided good descriptions of volume conduction effects on recorded signals. For example, CSF has shunting effects on volume currents, either amplifying or attenuating the magnitude of the scalp potential (van den Broek et al., 1998; Vorwerk et al., 2014). However, it may admittedly be difficult to understand the direct or practical implications of certain phenomena described in the biophysics literature for researchers working outside of that field. The aim of the present article is to bring knowledge from the field of biophysics about the impact of volume conduction on scalp EEG within the reach of scholars working with scalp EEG data and neurological populations, and provide some suggestions to address the issues. Conversely, we aim to highlight the use cases and final applications of such biophysical models.

### 1.1. Rationale of the present study

In general, when comparing a group with a CSF-filled lesion (e.g., stroke) with a lesion-free group, the lesion group will have more CSF due to the CSF-filled cavity and will have a particular distribution of the CSF inside the head. This creates systematic anatomical differences, requiring cautious interpretation of the comparisons of electrophysiological signals between the groups. Correlations between EEG-derived measures and severity of the damage (or of the resulting impairment) may also be confounded, as severity of impairment is highly correlated with lesion size (e.g., Schiemanck et al., 2005). That is, larger lesions are related to worse outcomes, but they also contain more CSF.

In light of the above, one may need to look at the observed scalp-EEG (difference) patterns commonly reported in the literature from a different perspective. When comparing a brain lesion group versus a lesion-free group, or when relating scalp-EEG patterns (e.g., ERPs) to impairment severity, both the differences in amplitude over the scalp and the unusual scalp distribution of ERP components can, in principle, be explained by volume conduction effects.

Here, we investigated the effect of lesions on the scalp EEG via simulations. More specifically, we simulated the EEG scalp potentials for the same active source(s), but using different volume conduction models, i.e. the anatomical models that govern the volume currents, using the finite element method (Engwer et al., 2017; Marin et al., 1998; Piastra et al., 2018; Vorwerk et al., 2018; Wolters et al., 2004). To approximate the scenario of a comparison between a lesion and a lesion-free group, in our simulations, we compared an undamaged brain, that is the reference volume conduction model with the geometrical and conductive properties of the tissues in the head, with the same reference model while adding a realistic CSF-filled lesion taken from empirical data (i.e., from stroke). We varied the size of the lesion by gradually adding realistically shaped CSF geometries to the reference volume conduction model to study the effect of lesion size, in analogy to associations between scalp-EEG patterns and impairment/lesion severity.

We then simulated EEG scalp potentials for known sources by “injecting” the same known signal through those different volume conduction models (also known as the “forward solution” when doing “forward modelling”). We used this approach for 1) a scenario with a single dipole source (a proxy for early sensory components such as visual evoked potential, e.g., (Arroyo et al., 1997) about 6 mm (“close”) or about 35 mm (“far”) from the CSF-lesion cavity and 2) a scenario with a distributed configuration of sources (a proxy for a cognitive ERP such as the P300 or N400 components, e.g., Bledowski et al., 2004; Halgren et al., 2002).

Since the only difference between the brain activity being compared is the amount of CSF present, we can evaluate the effect the CSF-filled cavity has on signal magnitude and signal topography (in the absence of external, channel-level noise). This enables us to critically assess the extent to which scalp EEG findings in populations with brain damage resulting in CSF-filled cavities (e.g., stroke or resections) can be interpreted at the level of cognitive functions, or may also, rather more trivially, be partly explained by properties of the volume conductor.

## 2. Methods

A general overview of the analysis pipeline is presented in Figure 1. All scripts and data for this study can be found at https://data.ru.nl/login/reviewer-2693535800/F37GZIPKCAJQFXMCNRUTUJTNU4ZXAHWZBN6IPMQ.

**Figure 1.**
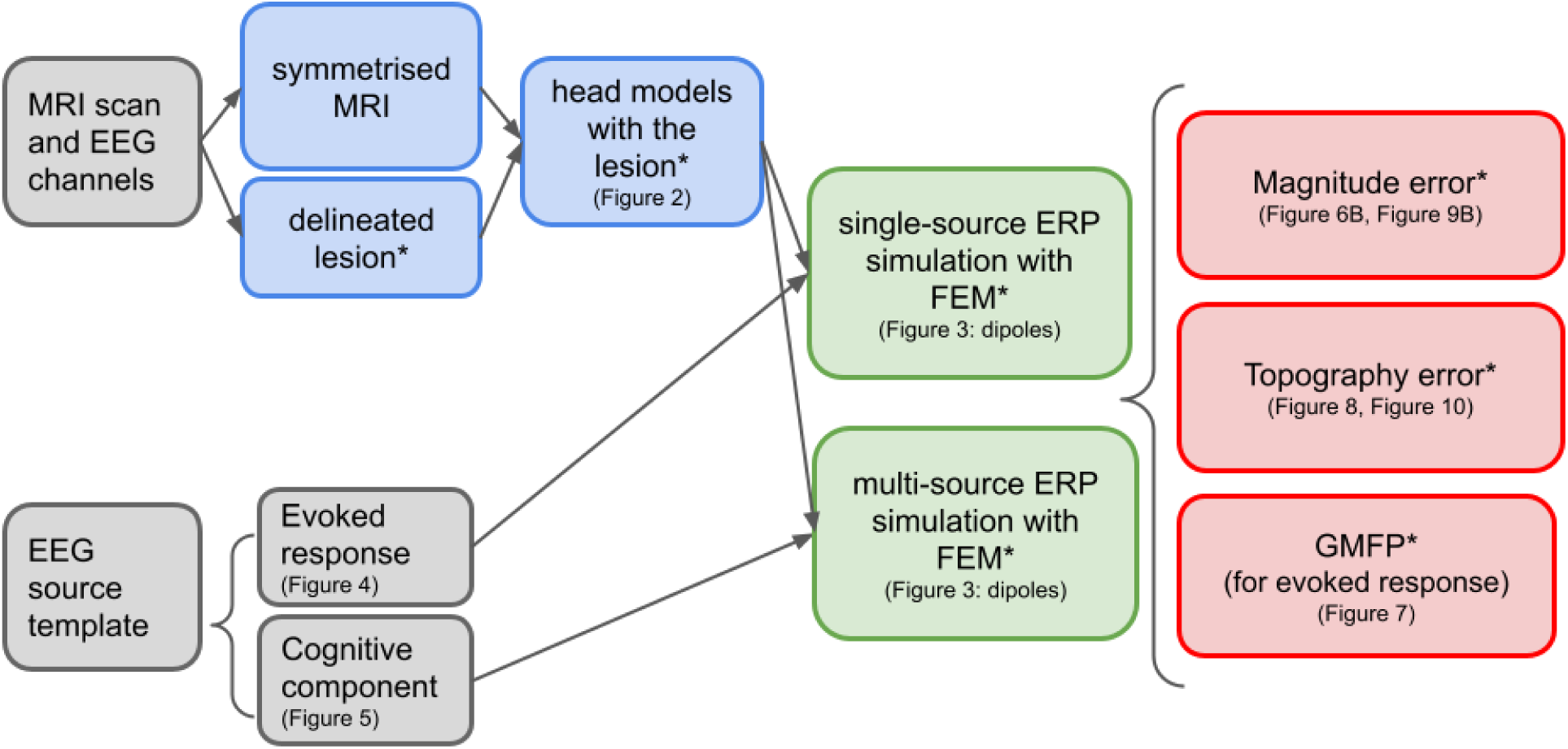
Analysis pipeline. ERP = event-related potential; FEM = finite element method; GMFP = global mean field power. Asterisks indicate the steps that are repeated for the 11 different head models.

### 2.1. Data acquisition and selection

We used the anatomical T1-weighted magnetic resonance image (MRI) of an individual who suffered a stroke more than two years before the scan, which was acquired under the approval of the Ethics Committee ‘CMO regio Arnhem-Nijmegen’, NL58437.091.17, following written informed consent. The participant was scanned at the Donders Centre for Cognitive Neuroimaging with a 3T MAGNETOM PrismaFit scanner. Realistic ERPs were derived from a study by Lewis et al. (2017) with visual stimulation and a study by Abrahamse et al. (2021) with auditory stimulation in an oddball paradigm (detailed below).

### 2.2. Modelling the head, the sources, and the electrodes

The MRI was symmetrised along the midline (explained in detail in Piastra et al., 2022) and segmented into five different components, using spm12 (Ashburner & Friston, 2005) with corresponding conductivity values assigned following the literature (grey matter = 0.33 S∕m, white matter = 0.14 S∕m; CSF = 1.79 S∕m; skull= 0.01 S∕m; scalp = 0.43 S∕m; Baumann et al., 1997; Dannhauer et al., 2011; Ramon et al., 2003).

This symmetrical head model (i.e., reference model) represents a participant without brain damage (i.e., 0% of the original lesion mask, left-most model in Figure 2). In the original MRI image, the lesion was delineated, and this delineated lesion was added as an additional CSF compartment, gradually extended in the anterior direction in steps of 10% until it corresponded to the total size of the original lesion (right-most model in Figure 2). This procedure resulted in ten additional head models with incrementally larger lesions, representing brain-damaged participants with varying lesion sizes. The smallest lesion (10% of the original lesion mask, second left-most model in Figure 2) comprised 4 ml, corresponding to 0.3% of the total brain (grey plus white matter) volume. The largest lesion (100% of the original lesion mask, right-most model in Figure 2) comprised 122 ml, corresponding to 9.4% of the total brain volume. The segmented volumes were downsampled to 2 mm resolution and used for building hexahedral volumetric meshes (with approximately 4.8 M nodes and 4.7 M hexahedrons).

**Figure 2.**
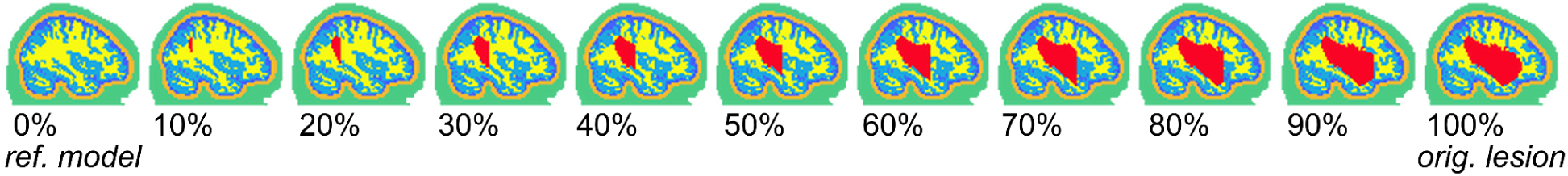
Segmentation (colour coded) of the 11 head models. Using the original lesion delineation and a symmetrical MRI, a lesion compartment was gradually introduced in increments of 10%. Segmentation indicates grey matter, white matter, CSF, skull, scalp, and lesion (in red). The lesion was modelled as CSF. Left hemisphere is shown. ref. = reference; orig. = original.

Sources were modelled as point-like equivalent current dipoles (de Munck et al., 1988; Murakami & Okada, 2006) in the grey matter compartment with an orientation perpendicular to the local cortical folding. The dipoles are visualised in Figure 3. There were two types of simulations: single source (Figure 3A) and multi source (Figure 3B). For the single-source ERP simulation, we conducted two forward simulations: one with the dipole positioned further away from the lesion in occipital cortex (at the shortest distance of 35 mm from the lesion), and one with the dipole positioned in the proximity of the lesion (at the shortest distance of 6 mm from the lesion). For the multi-source ERP simulation, a distributed configuration of dipoles was used. The shortest distances relative to the lesion surface were 15 mm, 8 mm, 10 mm, and 6 mm, respectively, for the dipoles in Figure 3B, going from the left-most to the right-most dipole anti-clockwise.

**Figure 3.**
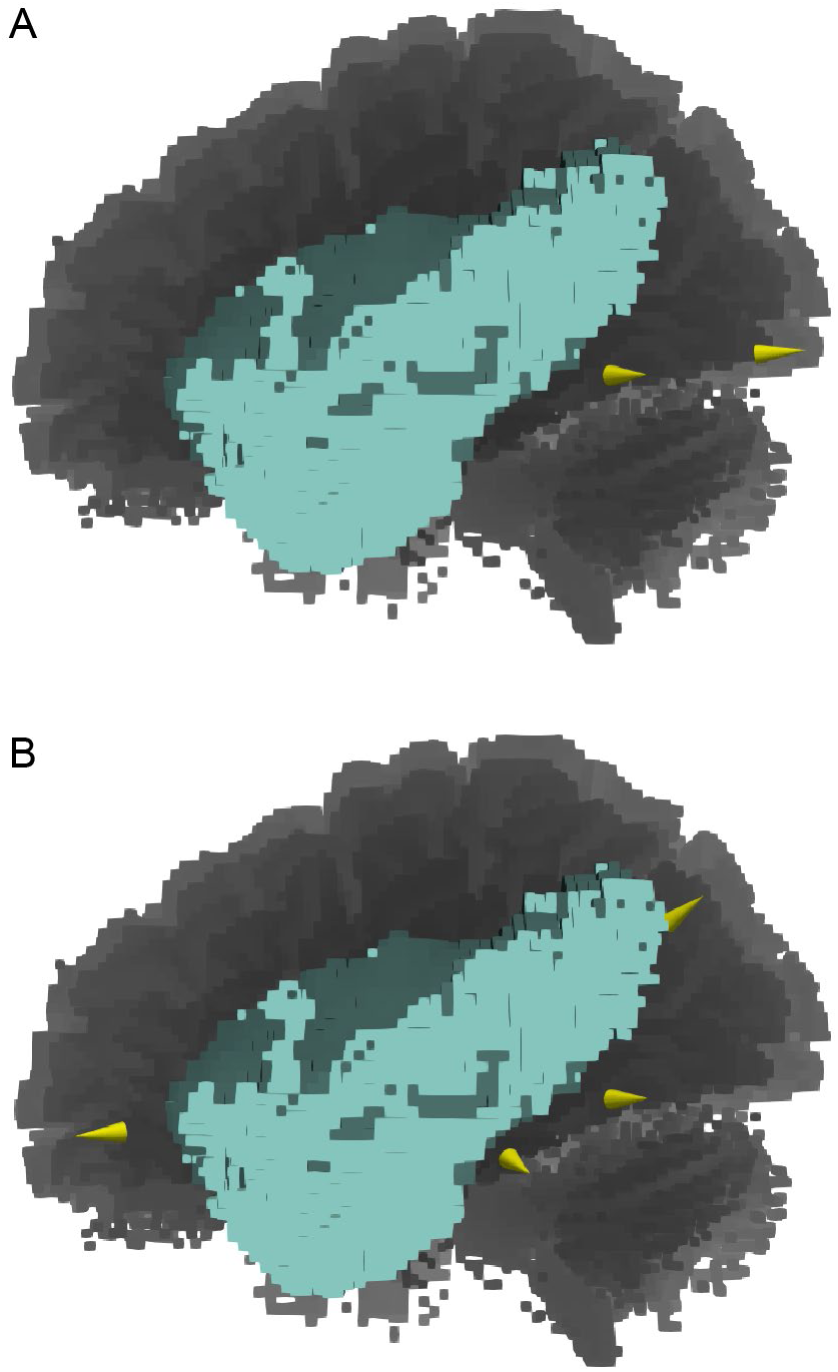
Dipoles for the single-source ERP simulations (A) for the source closer to the lesion (left-most dipole) and further away from the lesion (right-most dipole), and for the multi-source ERP simulation with a distributed configuration of dipoles (B). Blue shades: lesion; grey shades: grey and white matter; yellow: dipoles.

Template EEG electrode positions were used following the 10-20 system for 64 electrodes. The electrodes were aligned to the MRI using three fiducial points (nasion, and left and right preauricular points).

### 2.3. EEG forward solutions

We computed the EEG forward solutions for the different dipole configurations, applying the finite element method (FEM) implemented in FieldTrip (based on the SimBio software, Vorwerk et al., 2018) using the 11 different volume conduction models (1 non-lesioned, 10 lesioned). Rather than the instantaneous brain state, here, we were interested in studying the dynamics of brain states, i.e., stimulus-evoked (event-related) potentials. For this, scalp EEG signals were generated by combining the forward solutions with simulated activity time courses of the dipoles involved. That is, although the FEM solution is static over time, we obtained the dynamics of brain states by multiplying this FEM solution with the activity time courses of the source(s) involved. These activity time courses were derived from real data. Specifically, we used EEG data of a visual stimulation experiment for the single-source ERP simulation, and EEG data of an auditory oddball experiment for the multi-source ERP simulation, as explained below.

#### 2.3.1. Single-source ERP simulation

To simulate an early sensory component, such as a visual evoked potential, we assume a specific activity time course that is generated in a cortical patch comprising a group of pyramidal cells (i.e., the dipole). In our simulations, we positioned the dipole in two different locations: further away or closer to the lesion, and assigned an orientation perpendicular to the local cortical folding (i.e., a radial dipole on the apex of the gyrus). For the sake of comparability, the time course of activity at the two locations is assumed to be identical (see Figure 4).

**Figure 4.**
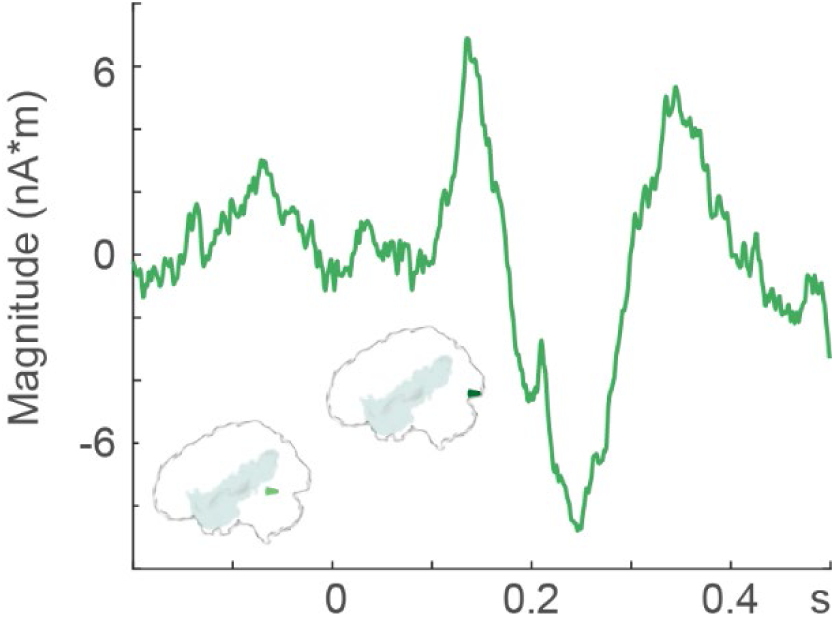
Single-source activity time course and dipole closer to the lesion (light green) and further away from the lesion (dark green). The same time course of activity was used for both dipoles.

The specific activity time course was constructed from an existing data set with visual stimulation (Lewis et al., 2017). Artefact-free scalp EEG data was segmented between –0.2 s to 0.5 s relative to the onset of a visual stimulus and averaged over channels ‘Oz’, ‘O1’, and ‘O2’ and averaged over 98 artefact-free trials, yielding an averaged VEP response that we assume to represent the cortical activity.

This activity time course was assigned to the simulated source and projected to the scalp for all 11 head models with the dipole placed in occipital cortex (35 mm from the lesion) and again for all 11 head models with the dipole in posterior proximity to the lesion (6 mm from the lesion). Figure 4 shows the dipole activity time course.

#### 2.3.2. Multi-source ERP simulation

Here, we were interested in cognitive event-related potentials, such as the deviant and the standard in an auditory oddball paradigm, or semantic violations and regular sentences in a language task. These ERPs measured at the scalp (and their associated components, such as the P300 or N400 effects) are known to be generated by a distributed configuration of sources (Bledowski et al., 2004; Halgren et al., 2002; Jentzsch & Sommer, 2001; Lau et al., 2008). We created a scenario where we have four different cortical patches (i.e., four dipoles) in different locations distributed within one hemisphere, each one with its respective standard and deviant activity time course. All four dipoles have an orientation perpendicular to the local cortical folding, on the apex of a gyrus.

The specific activity time courses were generated based on an existing auditory oddball paradigm dataset (Abrahamse et al., 2021), with a deviant and a standard condition, in the following way. Artefact-free scalp EEG data was segmented between –300 ms to 1000 ms relative to auditory stimulus onset. Independent component analysis (ICA) was used to decompose the signal. The first four components (comprising both standard and deviant trials) were assumed to represent the activity time courses from four cortical sources. Per experimental condition, we averaged each component’s time courses over trials (362 standard and 57 deviant trials), creating four specific single-dipole activity time courses per experimental condition. The deviant and the standard conditions will henceforth be referred to as condition 1 and condition 2, respectively.

Four different dipole locations around the lesion were chosen, representing a potential neuronal configuration of a cognitive ERP, and each dipole location was assigned one set of condition 1 and condition 2 dipole activity time courses. These dipoles are shown in Figure 5 with their corresponding activity time courses. Finally, the individual signals projected to the scalp with the FEM solution from these four separate dipoles per condition were summed at the EEG scalp level, resulting in one simulated condition 1 ERP and one simulated condition 2 ERP.

**Figure 5.**
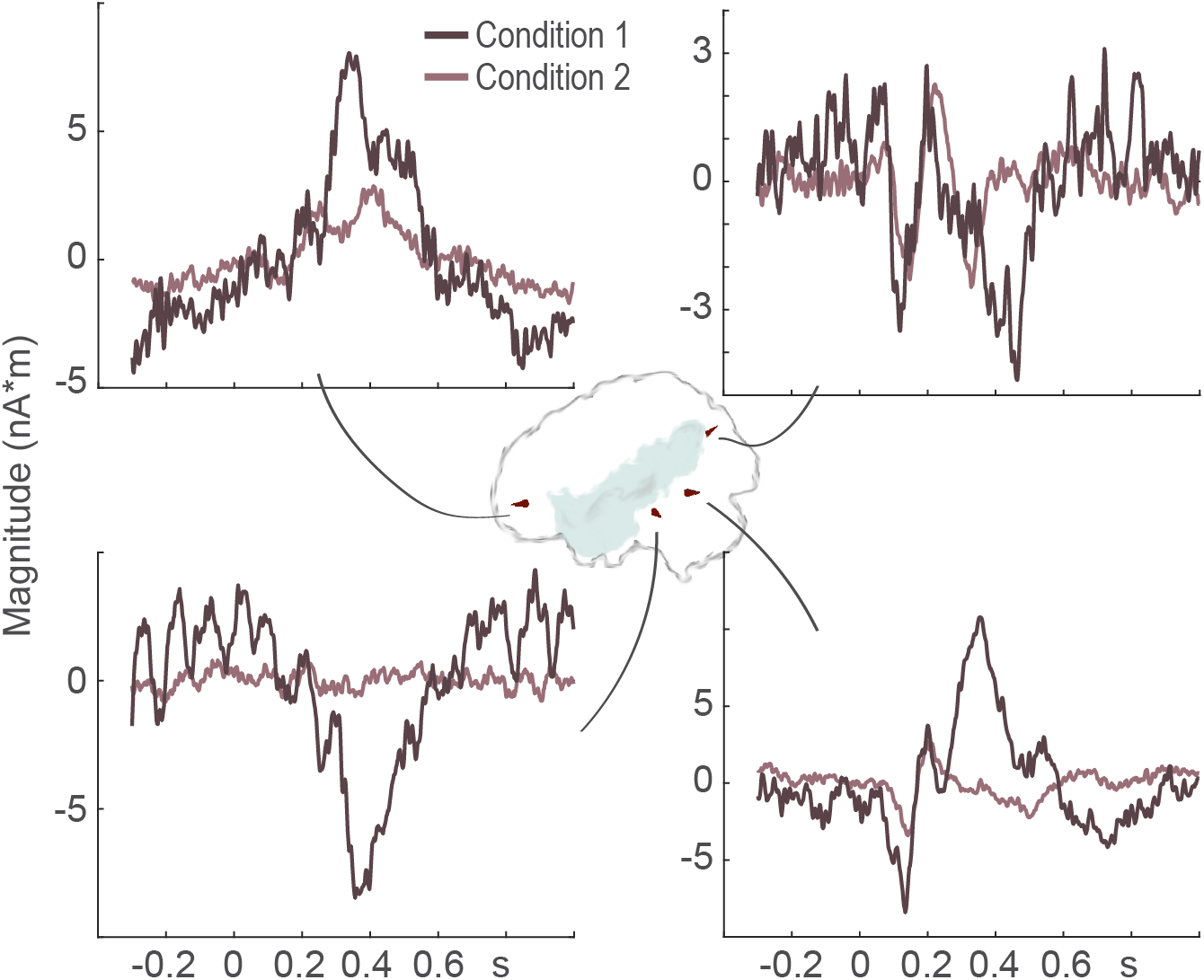
Time courses of the activity for four sources in a cognitive paradigm with condition 1 and condition 2 (taken from an empirical P300 oddball task). Each set of condition 1 and condition 2 dipole activity time courses was assigned to one dipole, as indicated.

### 2.4. Evaluating the effect of omitting the lesion on scalp-level interpretation

We examined the simulated EEG signal at the scalp level, quantifying the error in magnitude and topography relative to the reference model (i.e., 0% model; Meijs et al., 1989; Piastra et al., 2018) as a function of lesion size.

Magnitude differences between the reference model (i.e., 0% model) and the lesion models were quantified by the magnitude difference measure in percentage (MAG%), after averaging the signal within the relevant time windows defined in detail below, using formula (1).

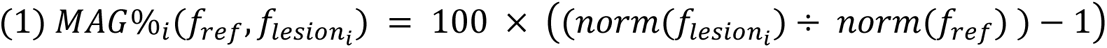

Differences in topography as a function of lesion size were quantified between the reference model and the lesion models using the relative difference measure in terms of percentage (RDM%) for the same signal and time window following formula (2):

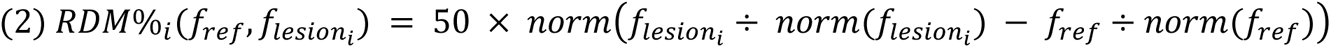

In both equation 1 and 2, *f_ref_* is the vector of the mean ERP amplitude within the time window of interest for all 64 electrodes for the reference model and *f_lesion_* the vector of the mean ERP amplitude over the same time window for all 64 electrodes for each *i* of the ten lesioned models. Values closer to zero indicate smaller differences in signal magnitude and topography.

#### 2.4.1. Single-source ERP simulation

For this simulation, the time window of interest was between 210 to 270 ms (i.e., when the negative-going deflection in the original dipole signal is largest, see Figure 4). Additionally, given that the Global Mean Field Power (GMFP) is used in the literature to characterise global EEG activity, we calculated the GMFP of the forward simulated signals in all 11 models using the formula from Esser et al. (2006). The GMFP was used here for visualisation purposes only.

#### 2.4.2. Multi-source ERP simulation

For this simulation, firstly, an ERP difference wave was calculated (condition 1 minus condition 2, the cognitive component/effect henceforth). Then, similarly to the single-source simulations, we computed MAG% and RDM% as a function of lesion size after averaging the cognitive ERP component (i.e., the difference wave) between 260 and 420 ms relative to stimulus onset (i.e., when the difference in deflection between the two conditions in the original dipole signal is largest, in line with the P300 literature, see Figure 5).

## 3. Results

### 3.1. Single-source ERP simulation

Figure 6A shows the simulated ERPs for all 11 head models at two channels ipsilateral to the lesion (left, upper row, ‘P7’; second row, ‘O1’) and one channel contralateral to the lesion left, third row, ‘FC4’) for the dipole close to the lesion, and for the channel P7 ipsilateral to the lesion for the dipole further away from the lesion (right, top row). The zoomed in ERPs are shown row-wise respectively to the right (grey shading). Electrode position and dipole (column-wise) are indicated for each panel with the simulated ERPs.

**Figure 6.**
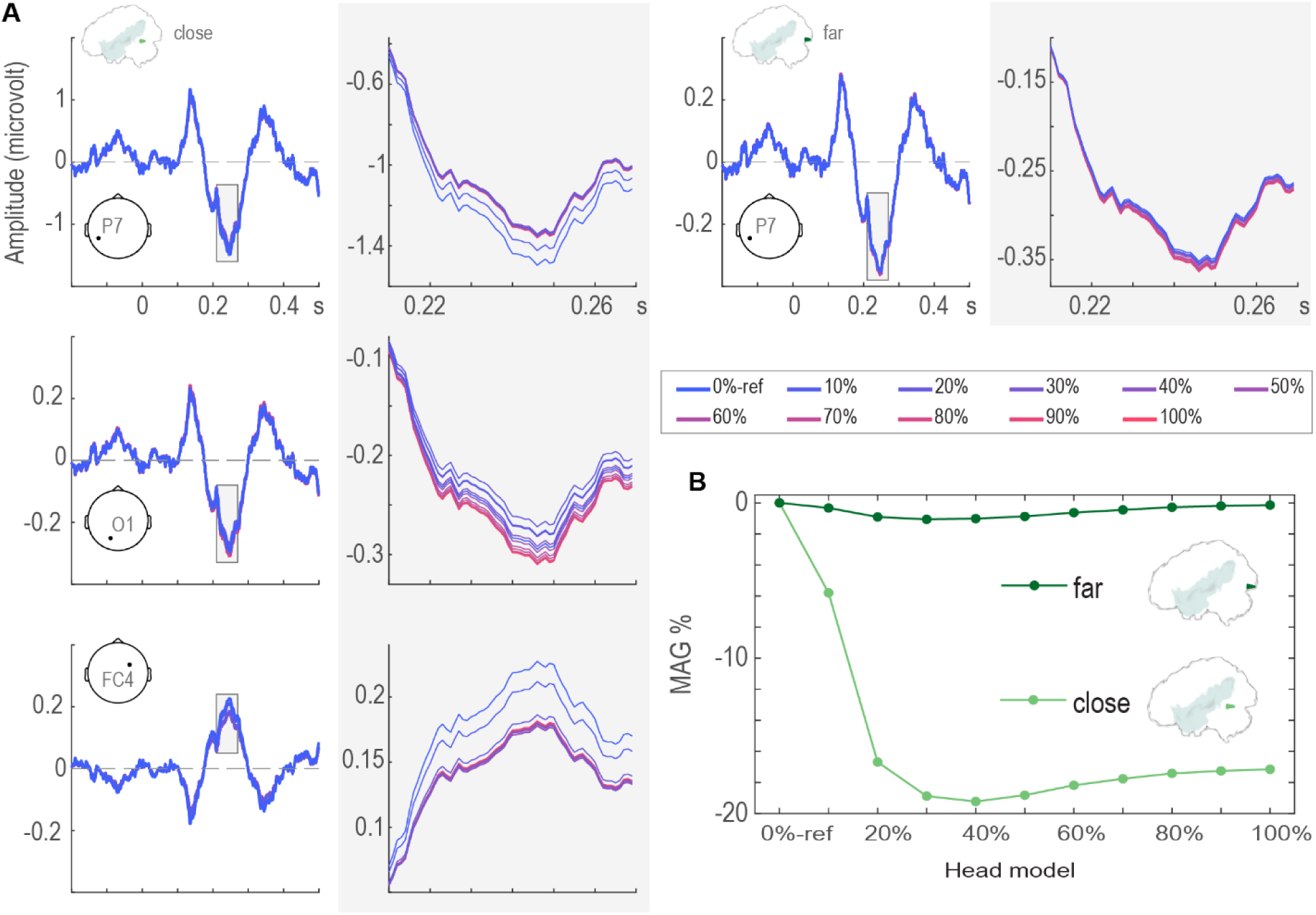
A. Scalp-level simulated evoked responses for all 11 head models for the dipole closer to the lesion at channel P7 (upper row, left) and for the dipole further from the lesion at channel P7 (upper row, right), as well as for the dipole closer to the lesion at channel O1 (second row) and FC4 (third row). Right-hand side panels with shaded background display the zoomed-in results between 210 and 270 ms (as indicated by the shaded area in the left-column plots) for each respective row. Coloured lines correspond to the 11 head models (right bottom box). Electrode positions are indicated for each panel with the evoked responses. Dipoles are indicated at the top of the columns. **B.** Magnitude difference measure in percent (MAG%) for magnitude differences between the reference model (0%-ref) and the lesion models as a function of lesion size for the dipole closer to the lesion (light green) and the dipole further away from the lesion (dark green). The MAG% between the reference model (0%-ref) and itself, which is by definition zero, is shown for completeness.

The amplitude at a same electrode site is modulated by the size of the CSF-filled compartment for the dipole closer to the lesion (three rows, left column), but not, or negligibly so, for the dipole further away from the lesion (top row, right column). Note that, for this reason, ERPs are shown for only one electrode for the dipole further away from the lesion. Figure 6B shows this phenomenon quantified in terms of MAG%. Note that the polarity of the input signal within the selected time window is negative. Larger magnitude error MAG% between the lesion models and the reference model is represented by more extreme negative values.

For the dipole close to the lesion (light green in Figure 6B), the mean ERP amplitude is related to lesion size in a somewhat monotonic fashion for increasing lesion size until the 40% model. The amplitude is attenuated to a similar extent (with more than 17% magnitude error) for all lesion models between 20% and 100% of the original lesion size. This effect is observed not only at an electrode on top of the simulated dipole (‘P7’, Figure 6A, top row in left column), but also at an electrode contralateral to the lesion (‘FC4’, Figure 6A, third row in left column). For the dipole further away from the lesion (Figure 6A, right column), lesion size has a very small effect on the scalp ERP amplitude, with attenuation values of less than 2% (dark green in Figure 6B).

Figure 7 shows the dependence of the GMFP on the lesion size. It is clear that the modulation as a function of lesion size observed in Figure 6 for the electrode-level ERPs for the dipole closer to the lesion is not amended by using the GMFP.

**Figure 7.**
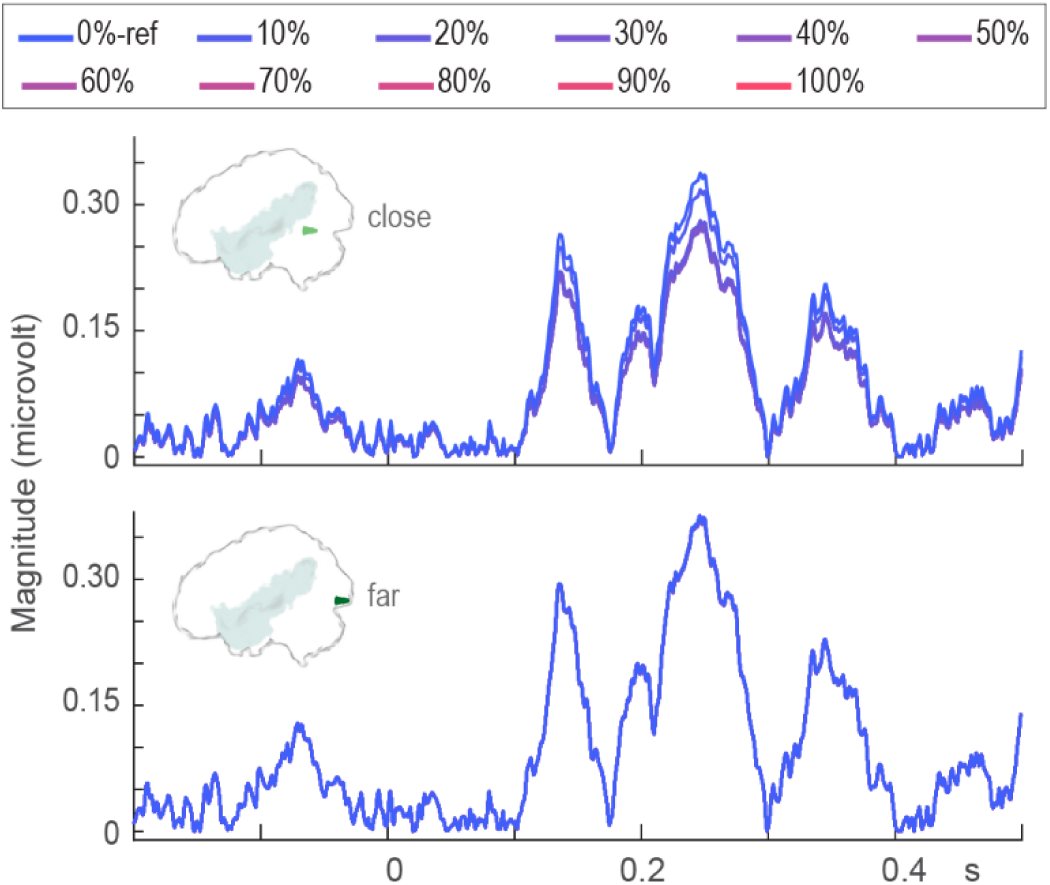
Global mean field power of the simulated evoked responses for both dipole locations as indicated in the panels (top, closer to the lesion; bottom, further away from the lesion) and all 11 head models. Coloured lines (top) correspond to the 11 head models. The dipoles are indicated.

Topographical changes of about 9% are observed for the dipole close to the lesion, starting from the 40% lesion model. For the dipole far from the lesion, the lesion size has little impact on the topography of the signal. The topography of the forward simulated ERP averaged between 210 and 270 ms (time window indicated by the shaded boxes in Figure 6A) for a selection of the head models (0%-reference, 20%, 100%) is shown in Figure 8 in blue-red colour scale. The *difference* in topography between the 0%-reference and the 20% and 100% head models is also shown in Figure 8, in purple-orange colour scale. The topography error for each of the lesion models relative to the reference model in terms of RDM% for the dipole close to the lesion (light green) and far from the lesion (dark green) are also shown in Figure 8.

**Figure 8.**
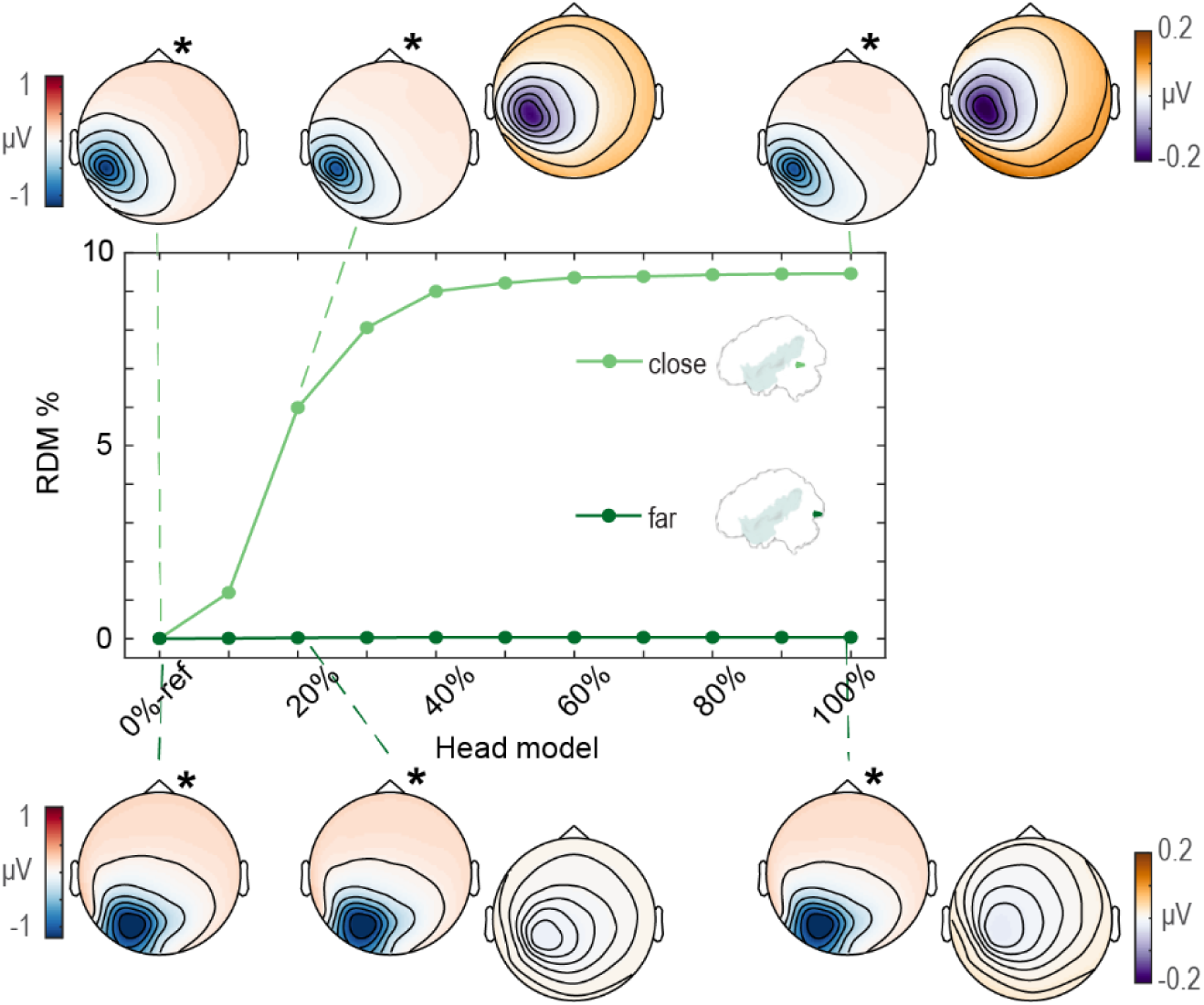
Relative difference measure in percentage (RDM%) for topography differences between the reference model (0%-ref) and the lesion models as a function of lesion size for the single-source simulations for the source closer to the lesion (light green) and the source further away from the lesion (dark green). Dipoles are indicated in the inset brains. RDM% was calculated for the signal averaged between 210 ms to 270 ms. The full topography is shown in red-blue colour scale for the 0%-ref and 20% and 100% lesion models, also indicated by the asterisks. The topographies in orange-purple colour scale show the *difference* between the reference model and the 20% and 100% lesion models. The RDM% between the reference model and itself, which is by definition zero, is shown for completeness.

### 3.2. Multi-source ERP simulation

Figure 9A shows the ERPs for condition 1 (darker colours) and condition 2 (lighter colours) for the 0%-reference (olive/sand) and 100% lesioned (wine/rose) models at four different electrodes for the distributed dipole configuration. As is clear from the figure, the amplitude at a same electrode site is modulated by the size of the CSF-filled compartment, especially in the case of condition 1, for which the amplitude is larger relative to condition 2. Figure 9A (shaded background) shows the zoomed-in cognitive effect (difference wave) within the interval of 260 and 420 ms post-stimulus onset for all 11 head models.

**Figure 9.**
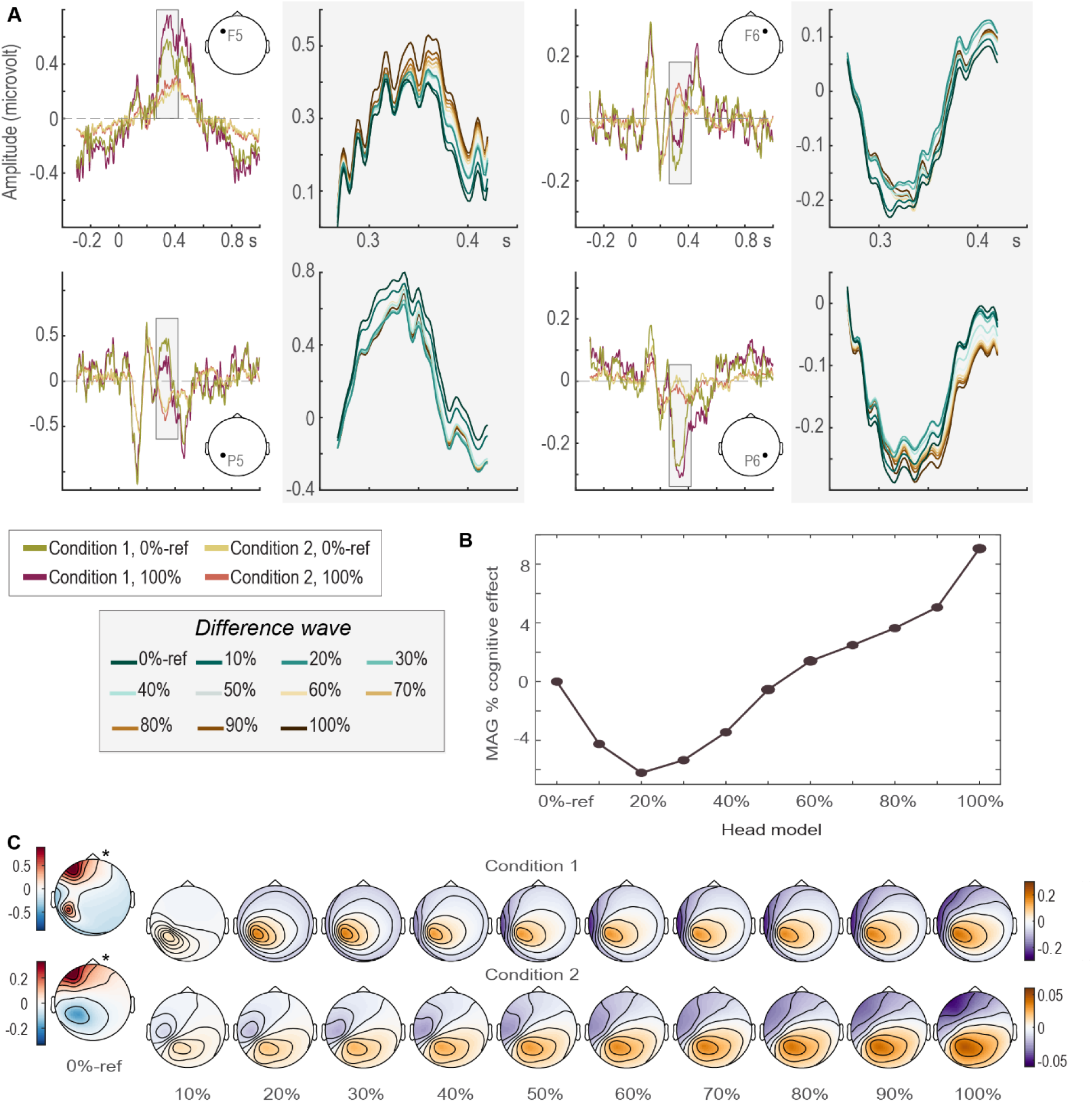
A. Scalp-level simulated ERP responses for condition 1 (darker colours) and condition 2 (light colours) for the 0%-reference model (olive/sand) and 100% lesioned model (wine/rose) at channel F5 (upper left), and F6 (upper right), P5 (lower left), and P6 (lower right), and corresponding zoomed-in ERPs (shaded background) to the right of each channel-specific panel. Zoomed-in results show the cognitive ERP effect (condition 1 minus condition 2) between 260 and 420 ms as indicated by the shaded areas for all head models. Colours corresponding to the 11 head models are shown in the bottom left box (shaded background). Electrode position is indicated for each panel. **B.** Magnitude difference in percent (MAG%) between the reference model (0%-ref) and the lesion models as a function of lesion size for the cognitive ERP effect averaged between 260 and 420 ms. The MAG% between the reference model and itself, which by definition is zero, is shown for completeness. **C.** Full topography of condition 1 (upper) and condition 2 (lower) averaged between 260 and 420 ms for the 0%-ref model (left, in red-blue, also indicated by the asterisks), and *difference* topographies between the 0%-ref model and all subsequent lesion models (orange-purple).

Figure 9B provides a summary of the effect of lesion size on the magnitude of the cognitive ERP effect (i.e., condition 1 minus condition 2) averaged between 260 and 420 ms post-stimulus onset in terms of MAG%. Changes in the amplitude of the cognitive effect as a function of lesion size show a complex, non-monotonic pattern, ranging between –6% to 9%, different from the pattern seen for the single-source simulation. Note that for the multi-source simulation, the lesion size gradually increases towards anterior sites in the left hemisphere, interacting in a complex way with the distance between the lesion (CSF) compartment and the different sources in the distributed dipole configuration.

The topography of the cognitive ERP effect averaged between 260 and 420 ms for each of the head models, as well as the *difference* between the 0%-reference and three of the ten lesion models is shown in Figure 10. The differences in topography for each of the lesion models relative to the reference model in terms of RDM% discloses again a non-monotonic relationship between lesion size and topographical distribution. Topographical changes between 16% and 18% are observed for all lesion models relative to the reference model, except for the smallest lesion of 10% (corresponding to 0.3% of the total grey plus white matter volume).

**Figure 10.**
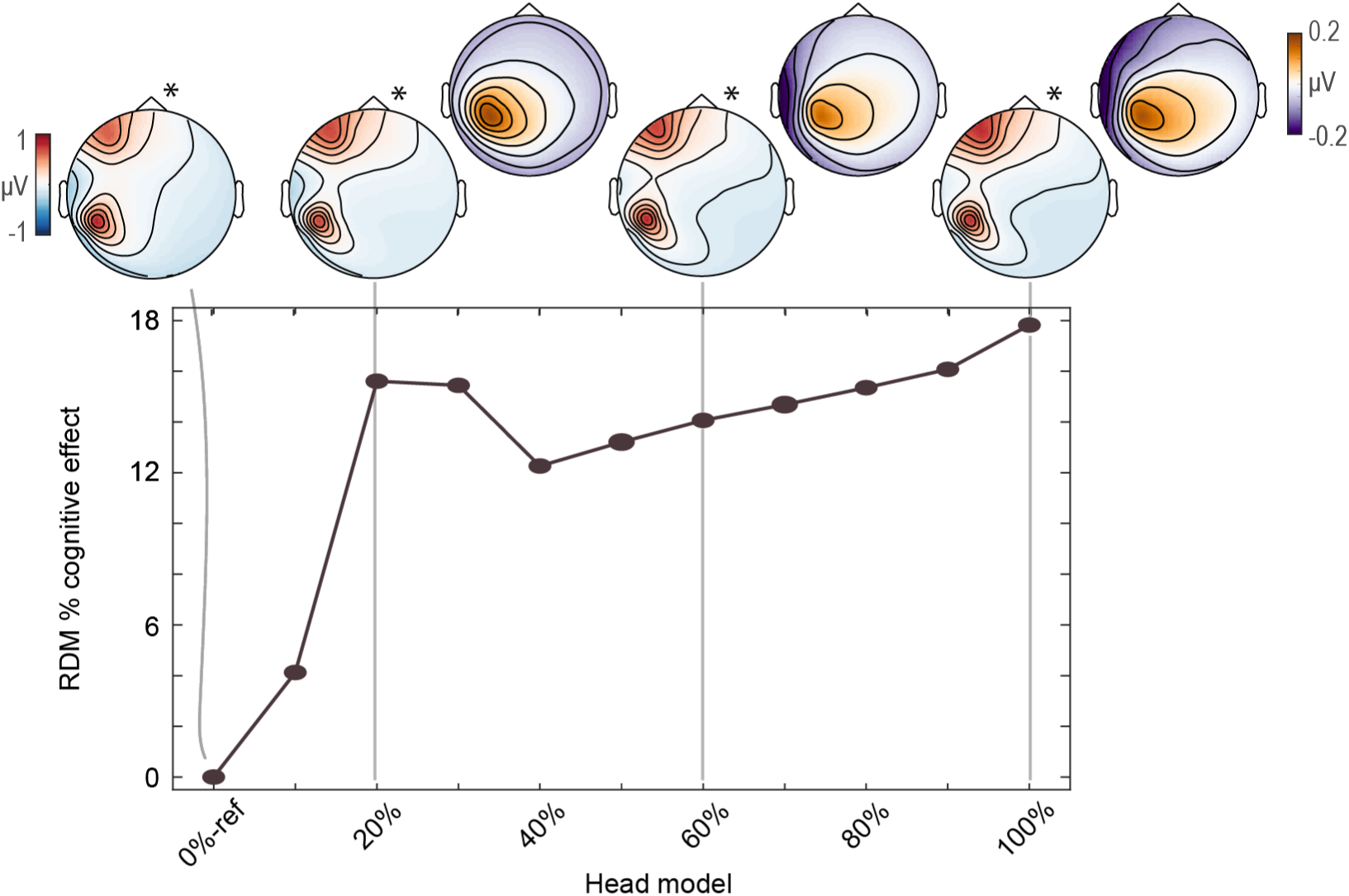
Relative difference measure in percentage (RDM%) for topography differences between the reference model (0%-ref) and the lesion models as a function of lesion size for the multi-source simulations for the cognitive effect (condition 1 minus condition 2) averaged between 260 and 420 ms. The RDM% between the reference model and itself, which by definition is zero, is shown for completeness. The full topography of the cognitive effect (averaged between 260 and 420 ms) is shown in red-blue colour scale for the 0%-ref and 20%, 60%, and 100% lesion models, also indicated by the asterisks. The topographies in orange-purple colour scale show the *difference* between the reference model and the 20%, 60%, and 100% lesion models.

## 4. Discussion

In the present study, we investigated the effect of CSF-filled cavities on the scalp EEG signal through forward simulations. We varied the size of the CSF-filled cavity spanning from the scenario of a brain without a lesion until a brain with a large lesion. We then injected the same signal through these different head models. We simulated a single-source scenario approximating an early sensory component being generated close to the lesion or further away from it, and a multi-source scenario approximating cognitive ERPs, for which a difference wave represents a cognitive effect. We reasoned that the only difference within each simulation scenario was the amount of CSF present, and its distribution, thus enabling us to critically assess the impact of the CSF-filled cavity onto scalp-EEG signal magnitude and topography in the absence of channel-level noise.

For the single-source scenario with the dipole located about 6 mm from the lesion, we found that the amplitude and topography of the signal at the scalp was affected by the size of the CSF-filled compartment, with amplitude attenuation of more than 16% for all but one lesion model, that is, for lesions larger than 12 ml. This attenuation was also observed when quantifying the signal in terms of the global mean field power. At the level of topographies, errors of about 9% are found for lesions of 37 ml or larger. For the dipole further away from the lesion, lesion size had a negligible effect on signal amplitude or topography.

For the multi-source scenario, we found that the amplitude and topography of the cognitive component (i.e., the difference wave) was modulated by the size of the CSF-filled compartment in a non-monotonic fashion, with up to 6% magnitude error (both attenuation and amplification) for relatively small and relatively large lesions, and more than 12% topography error for lesions larger than 12 ml. Our multi-source simulations indicate that focusing on an ERP effect, that is, taking a within-participant difference between two conditions and using that for comparisons of interest, as is done, for example, for the Mismatch Negativity, P300, or N400 components, does not remedy the issue.

Together, these findings have important implications for comparisons between a patient group with CSF-filled lesions and a lesion-free control group, as well as for relationships between patterns of the EEG signal and impairment severity. Even though our simulations are based on overly simplified scenarios in the absence of channel-level noise, the effects of the lesions are clear and often strong. We believe that our observations readily generalise to more realistic situations. Firstly, although very few, if any, sensorimotor or cognitive processes can be modelled with a single-source generator (e.g. early latency median nerve stimulation, e.g., Vanni et al., 1996), our single source simulations highlight the attenuation and topographical distortion of individual neuronal generators close to the lesions. It is safe to assume that sources underlying cognitive processes have more complex (often unknown) configurations, consisting of multiple dipolar sources with temporally overlapping activity time courses. Some of those individual sources might be close to the lesion, and the distortion of those components will have non-trivial effects on the mixed EEG-signals. For researchers performing lesion-based studies, the sources of interest may often be close to the lesion, thus requiring careful interpretation of any effects found. Secondly, our simulated lesions were relatively homogeneous and we gradually eroded the lesion in a very simplistic way (only in one direction), with the lesion shape changing predictably. By contrast, comparisons at the group level integrate across heterogeneous lesions. Whereas the impact of this predictable and gradual change was relatively monotonic in the single-source simulations, this was not the case in the multi-source simulation. The effect of lesion size and shape is complex, and no linear or monotonic outcome can be expected. This complex relationship between lesion size and geometry, on the one hand, and spatiotemporal characteristics of the active sources, on the other, means that simply adding lesion size as a covariate to a(n) often linear statistical model, as is common practice in the field, will not address the volume conduction confound.

Altogether, it is clear that for most of what cognitive neuroscientists would like to test, that is, multi-source configurations, the impact of a CSF-filled cavity cannot be neglected in the conclusions one wants to draw. Especially when comparisons are made at the group level, given the heterogeneity in size and shape of the lesions, any attenuated, aberrant, or absent effect at the scalp level will be very difficult to interpret.

Our simulations did not consider the influence of various sources of noise in great detail. Clearly, the reliability and validity of the measured signals are partly dependent on the signal-to-noise ratio (SNR). Here, signal is everything one is interested in measuring, whereas noise is everything one did not intend to measure. Put simply, in general, two types of noise sources can be considered, one internal and one external to the brain. One could argue that the effects we have shown on the signal magnitude as a function of lesion size are not likely to affect the signal’s SNR because the magnitude of both signal and noise would be amplified or attenuated due to the lesion, keeping the *ratio* constant. However, this view only considers internal sources of noise. External sources of noise (e.g., electrical interference, muscle activity, etc.) are not amplified or attenuated by the lesion. Given that the vast majority of the effects we observed on signal magnitude were one of attenuation, it is highly likely that the SNR will decrease as a function of the lesion. This fact impacts both comparisons between a lesioned and a non-lesioned control group, as well as correlations with impairment severity. Although we only demonstrated the impact of volume conduction on ERP patterns, the same outcome can be expected for other derived measures, such as time-frequency power, measures of connectivity, etc. This is because the volume conduction impacts the raw signal, thus affecting all measures computed from this raw signal.

This work provides some indication of when scalp patterns could be interpreted with less likelihood of volume conduction confounds. According to our simulations, lesions smaller than 1% of the total grey and white matter volume at least 35 mm away from the (single) source should not lead to dramatic changes to signal magnitude. As an example, this would represent a scenario of an investigation of early visual evoked responses (e.g., (Di Russo et al., 2005; Vanni et al., 2004) in individuals with frontal-lobe lesions. Critically, one needs to keep in mind that cognitive components tend to have a more complex and distributed configuration. The evaluation of within-patients measures of longitudinal changes over relatively short time periods (e.g. effects of language/cognitive/motor therapy) is likely less problematic. The reasoning behind this is that changes in CSF-related properties over time within a patient occur at a time scale that is different from the time scale at which the longitudinal measurements are compared. One would, however, ideally have to ascertain that the anatomy has not changed between the time points being compared, and that the SNR is comparable between the time points. With respect to the last point, for example, behavioural improvement over time may lead to a higher number of experimental trials to be acquired at a second time point relative to a first time point. This could already introduce SNR differences in the EEG signal between the comparisons, leading to a potentially erroneous conclusion that the signal reflects this behavioural improvement. Finally, the way to take the CSF-geometry issue into account is to perform analyses at the source level, where the effect of the CSF-filled cavity itself can be modelled (e.g., Piai et al., 2017; Piastra et al., 2022). Admittedly, this requires having detailed anatomical information available, including an anatomical scan, and technical expertise in source reconstruction, which are not always feasible scenarios.

On a final note, this work exemplifies use cases and final applications of biophysical models, which are often not the focus in more methodologically oriented studies. In cognitive studies in patient populations, for example, it is common to compare a brain lesion group versus a lesion-free group, or to relate scalp-EEG patterns (e.g., ERPs) to impairment severity, in particular by measuring both the differences in amplitude over the scalp and the unusual scalp distribution of ERP components. We hope that the present work can contribute to closing the gap between the methodological literature and final applications of these biophysical models.

### 4.1. Conclusions

Scalp-level comparisons between datasets with underlying differences in CSF anatomy do not always produce readily interpretable findings. Thus, comparisons between a patient group with and a control group without CSF-filled lesions as well as relationships between EEG scalp patterns and lesion size or impairment severity are likely problematic. The impact of the CSF-filled lesion extends beyond the amplitude of ERPs. Any measure dependent on magnitude or spatial patterns is potentially affected, as well as comparisons that, despite being somewhat independent on magnitude, will still be affected by differences in SNR. Lesion size in itself is not enough as a variable (to control for), as the most important feature is how the lesion relates to a source’s location and configuration. Although our results certainly do not mean that all effects found in the literature are caused by trivial volume conduction effects, claims about attenuated, aberrant, or absent effects at the scalp level based on comparisons with systematically different CSF geometry have to be made with caution.

## Acknowledgements

The authors would like to thank the Radboud University’s M/EEG community for input and, in particular, Ashley Lewis for kindly sharing data and Britta Westner for the insightful discussions.

